# A PAM-CNN recognition approach to Muscle Fatigue of Special Operators by sEMG signals

**DOI:** 10.1101/2025.07.20.665786

**Authors:** Xu Bin, Lin Minghui, Yue Zongzhen, Ji Shoucheng, Ouyang Weiping, Li Changyu

## Abstract

Fatigue operation was the primary accident causes of large special equipment which could lead to serious economic loss and personnel injury or death, so the establishment of early fatigue warning methods was of great significance to reduce the accident rate. In view of the working environment and biomechanics properties of operators of special equipment, theoretical and experimental method was presented to analyze the fatigue mechanism and accumulate data samples, and then a new approach was proposed to recognize muscle fatigue by classifying these datasets. Firstly, in order to deduce fatigue mechanism, hidden Markov chain was adopted to build five-element theoretical model for studying interaction between machines and operators’ muscles during operation process, so that influence factors and tested points of muscles were approached; Surface electromyographic (sEMG) sensors were introduced to acquire muscle signals which were taken as sample data, and wavelet packet was adopted to extract features of these data. Subsequently, a convolutional neural network (CNN) classification model based on partial attention mechanism (PAM) was proposed to execute triple fatigue classification, moreover cross entropy-Adam algorithm was applied to train and optimize this model. Finally, large crane operators were taken as case studies to verify the proposed model. By comparison results indicated when the number of extracted features was 3 and the size of patch was [5,5], PAM-CNN model could approach the optimum solution of fatigue recognition, and the average accuracy could achieve 95.6% in test datasets, which showed a good potential in practical application.

## 0 Introduction

Large special equipment was one of the essential infrastructure for development in manufacture and construction industry. Due to its extreme working conditions, accidents could result in serious disaster such as huge property losses and even personnel injuries, so safety was always the significant challenge to large special equipment. Operations of these equipment required constant attention from the operator as known to be a cognitively demanding task^[1]^, and fatigue had been identified as a key constraint that hampers the judgment and concentration of equipment operators, potentially resulting in accidents^[2]^, where it could account for more than 40% among accidents of cranes^[3]^. Fatigue recognition of operators was becoming an urgent subject to reduce the accident rate and improve safety and reliability of special equipment.

Fatigue is a complex multi-dimensional phenomenon which relates to the physiology, psychology and others, especially fatigue, drowsiness and sleepiness are often synonymous^[4]^. It is possible to distinguish among neural fatigue, fatigue of the neuromuscular junction, or muscle fatigue^[5]^. Muscle fatigue characterized by a reversible decline in the neuromuscular system’s force or power generation ability, can be categorized as central fatigue and peripheral fatigue^[6]^. Special operators such as crane operators should focused on muscle fatigue because of the decline of myogenetic vitality resulted from their long periodical and high load operation. The understanding and recognition of muscle fatigue have always been an unresolved topic and attract much more attention^[7]^.

As reviewed in the literature, electromyographic (EMG) signals had become an effective method in the field of muscle fatigue research, and non-invasive surface electromyographic (sEMG) sensors were more widely used because they did not require a needle-type puncture into the skin^[8]^. Jero^[9]^ used geometric features of sEMG to differentiate the muscle nonfatigue and fatigue conditions in isometric contractions. Greco^[10]^ divided 32 healthy subjects performing isometric biceps contraction into fatigued and non-fatigued group based on standard sEMG analysis. Reyna^[11]^ detected muscle fatigue in the shoulder and forearm caused by repetitive and continuous strain associated with the computer mouse using sEMG data. Zhao^[12]^ proposed a fatigue analysis method that fuses the improved EMG fatigue threshold algorithm with biomechanical analysis. Li^[13]^ proposed nine fatigue indicators derived from sEMG signals and a K-Nearest Neighbors (KNN) model was used predict muscular fatigue. CORVINI^[14]^ based on autoregressive model for muscle fatigue state identification through the mean and median frequencies of EMG signals. Xie^[15]^ assess the effect of self-contained breathing apparatus on muscle fatigue and compensation in firefighters using sEMG from the upper trunk muscles. Lin^[16]^ investigated the ability of turns-amplitude analysis and power spectral analysis of sEMG to identify fatigue and recovery under dynamic muscle contractions. These studies have applied various time-domain, frequency-domain, and complexity metrics for muscle fatigue assessment, and characterized motor unit (MU) features associated with muscle fatigue using high-densitys EMG^[17]^. Moreover, multi-modal sensors^[18]^ and multi-muscle synergistic effect^[19]^ were introduced to improve the detection accuracy.

Along with the great progress of artificial intelligence, machine learning had been widely used for sEMG signal process. The first step was feature extraction by algorithms such as Fourier transform, continuous wavelet transform^[20-22]^, etc. and then, machine learning algorithm was employed to realize fatigue recognition^[23-24]^. XU^[25]^ used multi-class Gaussian classifiers based on the time-frequency domain features of the sEMG for the muscle fatigue state identification. Zhang^[26]^ use the sEMG signal to detect muscle fatigue based on the Multidimensional Feature Fusion Network (MFFNet), which is composed of Attention Frequency domain Network (AFNet) and Attention Time domain Network (ATNet). Wang^[27]^ propose a neural architecture search framework based on reinforcement learning to autogenerate neural networks. Xia^[28]^ introduced multifractal detrended fluctuation analysis to extract non-linear properties of sEMG, and built a LSTM networks under the combined feature set. Temel^[29]^ applied artificial neural network and regression analysis for diagnosis of bruxism by features extracted from EMG signals. Sun^[30]^ used machine learning to classify muscle fatigue levels through EMG signals during isometric contraction.

The above-mentioned researches showed that it could achieve good accuracy to recognize muscle fatigue by sEMG signals. However, muscle fatigue is a nonlinear and non-stationary random process, and the classification model still needed to be further optimized in terms of self-learning ability and reliability. Based on attention mechanism (AM) and convolutional neural network (CNN), a Partial Attention Mechanism-Convolutional Neural Network (PAM-CNN) model was proposed in this study, and as a case study it was applied in fatigue recognition of large crane operators.

## 1. Mechanism of muscle fatigue

Large crane run in reciprocating motion for 24 hours continuously during their whole life cycle, which was manipulated mostly by operators other than automated control. Operators usually worked by shifts in eight hours one day and under the extreme conditions such as high altitude, limited space, hard environment and high intensity workloads. While operators manipulated the crane, muscles of arms and legs required quickly response to reciprocate motion, and these long-term repetitive monotonous movements would result in performance degradation, which was prone to muscle fatigue.

In order to reveal the mechanism of muscle fatigue and locate the muscles that appear fatigue firstly, theoretical analysis based on multi-body dynamics was introduced, the interaction between muscles and crane manipulators could be expressed as follows:

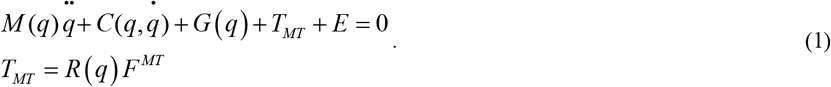

Where *q* was the angle of the *i*th biological joint, *M*(*q*) was the mass matrix of the system, 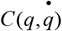 was the Cowlis load matrix, *G*(*q*) was the gravity load matrix, *T*_*MT*_ was the muscle joint moment matrix, *R*(*q*) was the force arm matrix of the muscle moment, *F*^MT^ was the muscle force matrix, and *E* was the external force (resistance during operation).

According to Hill rules, *F*^MT^ was affected by the number of muscle fibers, the average length of muscle fibers *L*, muscle mass *m*, muscle density *ρ*, and muscle strength coefficient *λ*, and its maximum value could be expressed as:

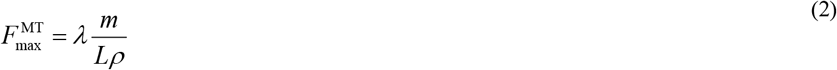

Where *λ* was the variable changing with time *λ*(*t*) during a certain period, so that 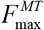 was transformed into a function varies with time 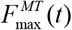

From equation(1)-(2), it indicated parameters of muscle forces and crane manipulators were difficult to achieve by experiments, and parameters in equation (2) of different muscles changed greatly. In addition, dynamic capacity of muscles changed with time and influenced the operation accuracy, so that it affected the dynamic properties of cranes. Simultaneously, the reaction forces exerted by the operating lever and pedal of cranes, coupled with the vibrations generated by cranes during operation, stimulated an interactive influence on muscles, which could manifest as a periodic human-machine coupling effect. Therefore analytical solution was not accurate to interpret the process of muscle fatigue of crane operators. Nonlinear hidden Markov chain model (NL-HMM) was proposed to approximate this random process, and sEMG signals was adopted as statistical data.

Since no definite mathematical equation could express the human-machine coupling effect until now, response surface method was adopted to fit this model. The response surface function was denote by *α*_*i*_ (*x*_*j*_) between muscle force 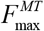 and operating parameters of the operating rod and pedal and kinematic parameters of cranes, and the response surface function was denoted by *β*_*j*_ (*y*_*i*_) between the reaction force and vibrations of the operating rod and pedal and features of muscle fatigue, and the interaction function was denoted by *δ*(*x* _*j*_ *y*_*i*_) between muscles and machines. NL-HMM was adopted to fit *β*_*j*_ (*y*_*i*_), with *p* denoting the set of muscle states, and *q* denoting the set of all possible observations, that is sEMG data.

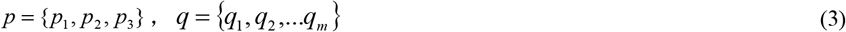

Where *p*_1_, *p*_2_, and *p*_3_ represented muscle normal, mild fatigue, and severe fatigue states; *m* represented the possible number of observations.

Denote the state transfer probability matrix by *A*

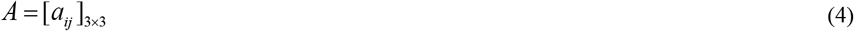

In matrix *A*,

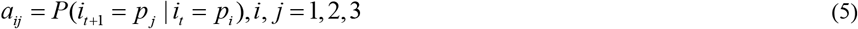

Denote the observation probability matrix by *B*,

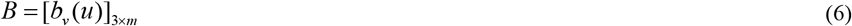

In matrix *B*,

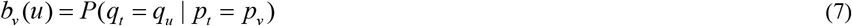

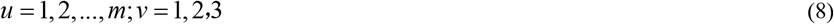

The probability vector *π* = (*π*_*y*_) represented the initial state with *π*, so the expression of *π*_*y*_ was:

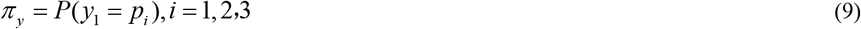

Based on the HMM model, a five-element model of muscle fatigue was conducted as:

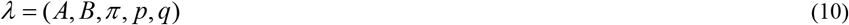

Fitting with the Chapman-Kolmogorol formula, the response surface model of the interaction *δ*(*x* _*j*_ *y*_*i*_) was established as:

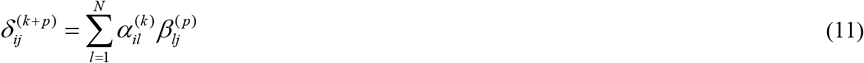

The time-series integration process of muscle forces was transformed into a discrete summation process. Above all, *P* in equation (5) was originally estimated by the working time, and *P* in equation (7) was originally provided according to empirical data. Therefore, there were many constraints of this analysis, in order to improve the precision a new machine learning model was proposed to optimized these parameters in the next step.

## 2. PAM-CNN model

Muscle fatigue is a non-stable stochastic process with Markov characteristics. It is difficult to obtain the characteristic information with high degree of separation just by time domain and frequency domain method, and the robustness of feature classification could not be guaranteed. Wavelet packet transform (WPT) was good at local feature extraction since it could characterize frequency and time features at different scales, and achieve time-varying spectral analysis, moreover it was insensitive to noise. Therefore, WPT method was used to carry out feature extraction of sEMG signals.

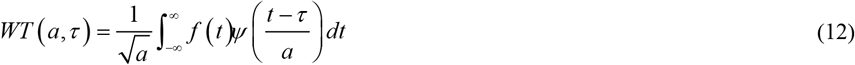

Where *a* was the scale of the wavelet function, *τ:* was time, *Ψ* was the wavelet function, and *f* (*t*) was the sEMG signal. After wavelet transform extraction, the sEMG signal was transformed into a two-dimensional signal containing time and frequency variables.

Introducing the ‘cmor’ wavelet as a wavelet basis function,

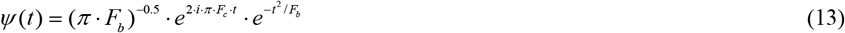

Where *F*_*b*_ was the bandwidth factor and *F*_*c*_ was the central frequency factor.

The features extracted by WPT were used as data samples for fatigue recognition, where labeled dataset was applied to train the classification model.

Since CNN had excellent autonomous learning capacity, it was widely used in pattern recognition. CNN used the gradient gradient descent method to minimize the loss function and retroegulate the weight parameters in network layer by layer. Two-dimensional convolution operation was adopted according to the scale of tested datasets and was described as follows:

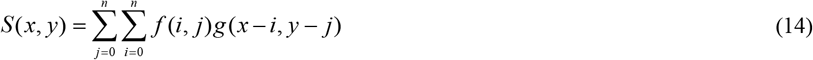

Where *f* (*i, j*) represented the input two-dimensional data, and *g*(*x* − *i, y* − *j*) represented the two-dimensional convolution kernel.

Since it needed lots of iterative training in the entire learning process, the computational efficiency was affected. The introduction of self-AM reduced iteration times significantly,

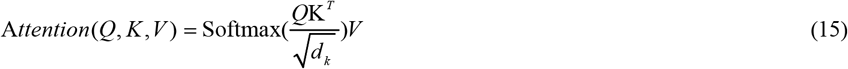

Where *Q* was the query set and *K* was the input data. The similarity information in each position of the image was obtained after the inner product of *Q* and *K*. Then the matrix of similarity coefficients was obtained after softmax normalization, and through the inner product with *V*, the important properties of the image information were achieved.

Since the samples received by sEMG sensors were very large, it resulted in a large feature subset. In order to increase training efficiency, PAM was proposed to apply the Spearman correlation analysis to rank the feature subset in terms of importance weights, and focus only on the fatigue-related feature subsets with the top ranks. *WT*_*Q*_ and *WT*_*K*_ were obtained from WPT processing of sEMG signals in fatigue state and normal state respectively, and the WPT sEMG signals were divided into multiple patch blocks.

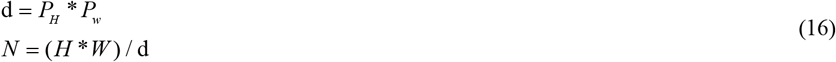

Where *d* was the size of the patch block, *P*_*H*_ and *P*_*w*_ were the height and width of the patch, *H* and *W* were the time axis and wavelet scale axis of sEMG signal after WPT processing, and *N* was the number of patch blocks obtained after segmentation.

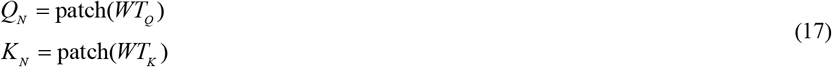

Where *Q*_*N*_ was used as the query set to segment sEMG signals in fatigue state, and *K*_*N*_ was the sEMG signal patch in normal state. Based on the principle of vector cosine similarity, the fatigue feature in the sEMG signal processed by WPT was obtained, that is

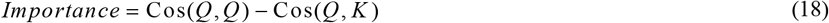

The importance weights were sorted into patch blocks, and Spearman correlation analysis was used to select the highly weighted and independent *Q*_*i*_ as the AM convolution kernel. Discrete summation was performed according to formula (11) for feature extraction of the sEMG signal.

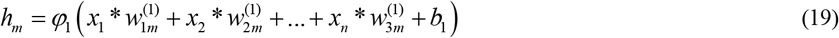

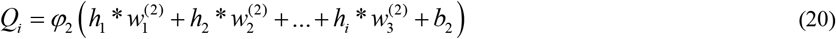

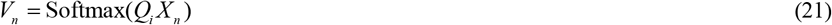

Where 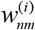 represented the weight of neurons between the perceptron layers, when *i* was equal to1, it represented the weight between the input layer and the hidden layer, and when *i* was equal to 2, it represented the weight between the hidden layer and the output layer. The weight value was calculated by the analytical model of muscle fatigue based on size of each sampled feature data.

Where *h*_*m*_ referred to the output of the *m* th neuron in the hidden layer, *x*_*n*_ referred to the output of the *n* th neuron in the input layer, φ_1_ was the activation function used in the hidden layer, and *b*_1_ was the bias input of the hidden layer. φ_2_ was the activation function used in the output layer, and *b*_2_ was the bias input of the output layer.

As a query array, fatigue features were extracted from the input sEMG signals *X*_*n*_, as shown in Figure 1. *V*_*n*_ was the fatigue information of patch *X*_*n*_ in *Q*_*i*_ at the time of *n*. Softmax function was applied to accomplish the normalization process and achieve the fatigue similarity information of *X*_*n*_.

**Figure 1.**
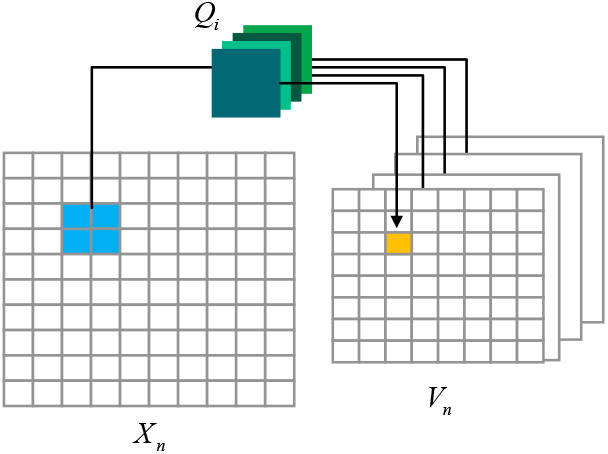
Feature extraction by PAM.

Fatigue features of sEMG signal was selected and optimized based on PAM model, and CNN was integrated to establish a PAM-CNN fatigue recognition model, as shown in Figure 2.

**Figure 2.**
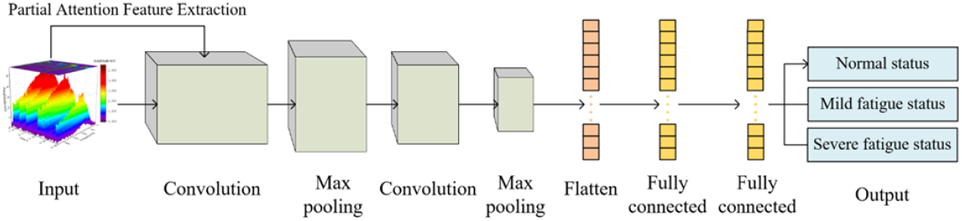
PAM-CNN model.

## 3. Case study

### 3.1 sEMG test

Container crane had the characteristics of large span, high lifting height, high operating frequency, and high full-load rate. It could lift 50-70 standard containers (Twenty-foot Equivalent Unit, TEU) per hour. Therefore, container crane operators were typical special operators and could be adopted as a standard sampling case for muscle fatigue process.

In order to reach the muscle fatigue quickly, 60 TEUs per hour were used as the tested case. During the operation cycle, movements such as operating the whole crane, the trolley and lifting machine were mainly completed by the left hand, while starting, stopping and emergency stopping of the electrical system were mainly completed by the right hand, and the acoustic and optical warning devices were generally operated with the right foot. Due to higher frequency of left-handed actions, samples of sEMG data from the left arm were chosen to be more representative.

Due to the significant differences in operating conditions of container cranes in different areas and periods, operators with similar working conditions in the same area were initially selected for sample collection. Considering the effect of lunch break on muscle fatigue, each shift from 9am to 11am was selected as a test cycle, and eleven crane operators with different age, height, and weight (as shown in Table 1) were chosen to establish the sEMG datasets of crane operators. Crane operators under normal state were tested without any exercises, while operators under fatigue state were tested after exercises with dumbbells, and data of exhausted exercises were labeled as severe fatigue state. The sEMG data of 9 operators were adopted as the training datasets, and the data of the remaining 2 operators were used as the test data, which was applied to validate the fatigue recognition model.

**Table 1.** Samples of crane operators

| Number | Gender | Age | Height<br>(cm) | Weight<br>(KG) | BMI index |
| --- | --- | --- | --- | --- | --- |
| 1 | male | 25 | 172 | 65 | 21.9 |
| 2 | male | 26 | 177 | 68 | 21.7 |
| 3 | male | 24 | 168 | 80 | 28.3 |
| 4 | male | 25 | 173 | 70 | 23.3 |
| 5 | male | 27 | 172 | 74 | 25.0 |
| 6 | male | 32 | 183 | 86 | 24.8 |
| 7 | male | 36 | 170 | 78 | 27.6 |
| 8 | male | 31 | 174 | 76 | 24.8 |
| 9 | male | 29 | 173 | 76 | 27.2 |
| 10 | male | 30 | 180 | 82 | 24.7 |
| 11 | male | 32 | 177 | 75 | 23.5 |

Each crane operator was monitored by sEMG sensors for a period of 5 days to obtain data samples for actual analysis. These tests were divided into twice considering the operating convenience and precision of sensors, one tested time was July17-21, 2023, and the other was May 13-17,2024. In addition, every participants provided informed consent by written documents. According to the characteristics of crane operation, and the dynamics state of muscles during operation, online monitoring of biceps and triceps muscles of the operator’s upper arm was carried out by a self-developed sEMG collector, which delay time was less than 500us, and sampling rate was 4,000 Hz. The sEMG electrode slices were distributed at positions as shown in Figure 3.

**Figure 3.**
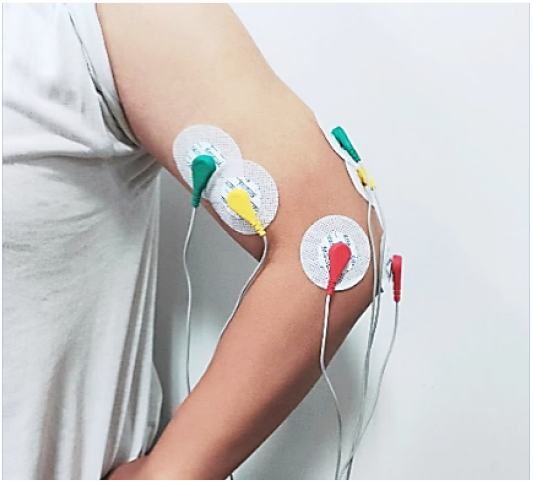
Positions of sEMG sensors.

The signals acquired by sEMG sensors were easily interfered by noise, and a narrow-band low-pass filter was adopted to filter out the high-frequency noise of the sEMG signals. Under the supervised mode, the validation set of PAM-CNN model was established by extracting the time domain features such as mean absolute value, number of zero-cross points, variance, root mean square, and frequency domain features such as power spectral density and average frequency of sEMG signal to label normal state, mild fatigue, and severe fatigue, as shown in Figure 4(a), 4(b), and 4(c).

**Figure 4.**
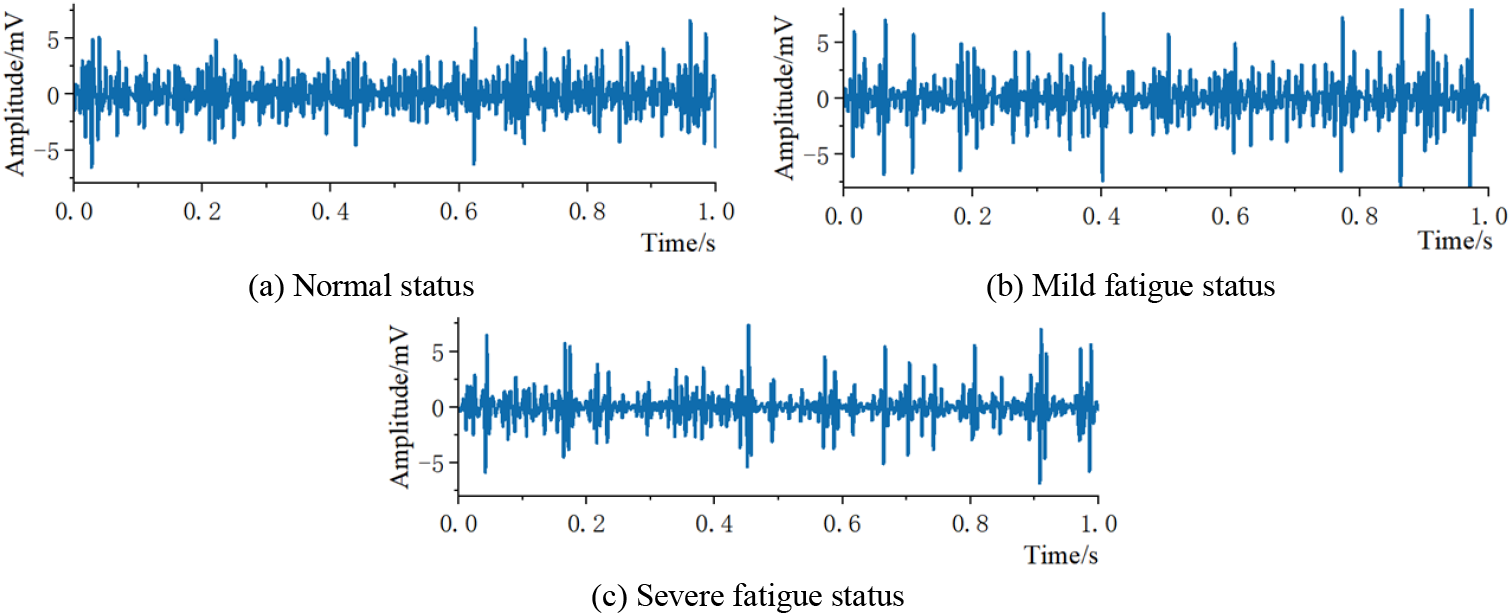
sEMG signals in time domain.

### 3.2 Feature extraction of sEMG signals

Before feature extraction, a time-overlapping sliding window was introduced to segment sEMG signal. The length of a single time window was set to 300ms, and the sliding interval between the windows was set to 200ms, so that the overlapping portion of the neighboring windows was 100ms as illustrated in Figure 5.

**Figure 5.**
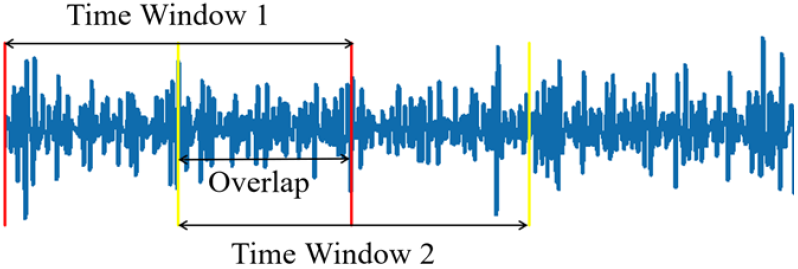
Segments of signals.

The cmor wavelet was introduced to extract features and the different fatigue states were shown in Figure 6. Different sizes of small windows [2,2], [3,3], [4,4], [5,5] were selected to slice the sEMG signals in 6(a) normal state and 6(b), 6(c) fatigue states. A query set Q was established based on the signal slices of fatigue state, and cosine similarity between the query set and slices in different states was obtained. According to equation (14), the slices of query set Q with high cosine similarity to the sEMG signal of fatigue state and low cosine similarity to the sEMG signal of normal state were selected as features of fatigue state.

**Figure 6.**
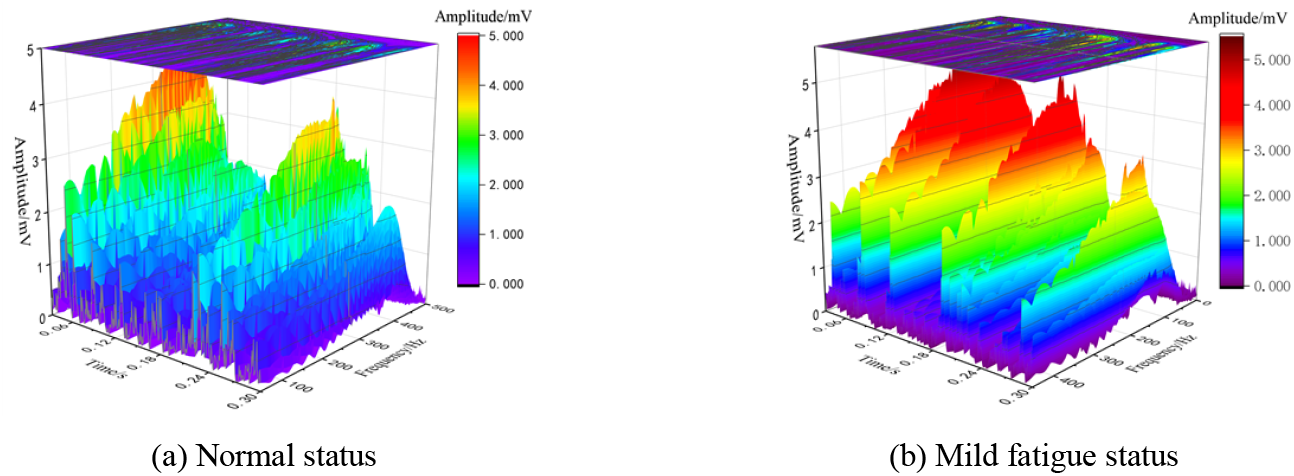
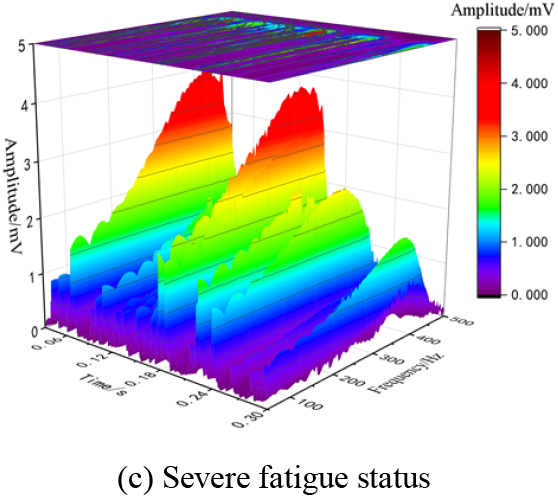
Time-frequency feature extraction of sEMG signals.

According to Spearman correlation analysis, independent fatigue features were obtained, and the extracted features at small window size [5,5] were shown in Figure 7, and the fatigue correlation of four sEMG features at each time period was shown in Figure 8.

**Figure 7.**
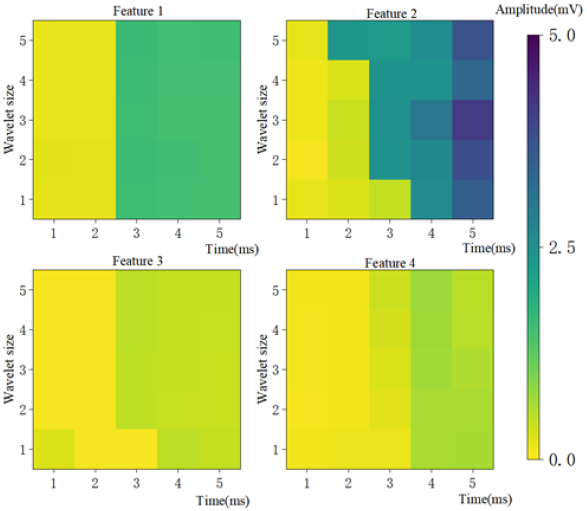
Four fatigue features splitted by small window.

**Figure 8.**
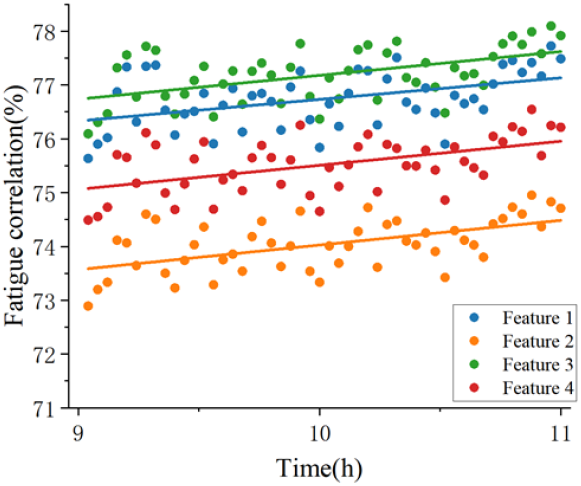
Fatigue relevance analysis.

### 3.3 Model training and optimization

Based on the PAM-CNN model, the sEMG features of tests were trained, and cross-entropy function was applied as the loss function, and adaptive moment estimation algorithm Adam was applied to the neural network back-propagation to iteratively optimize small window size and the quantity of features. The accuracy for extracting features with different small window sizes was shown in Figure 9, and its converged average accuracy and running speed were shown in Table 2. Taking the small window size [5,5] and training optimization for the number of extracted features, the accuracy of different number of features was shown in Figure 10, the average accuracy and running speed after convergence were shown in Table 3. When the number of iterations reached 150, the analytical accuracy of the training set and the test set reached the precision confidence interval, as shown in Figure 11.

**Table 2.** Model optimization results for different small window sizes

| Small window size | Running speed | Average accuracy |
| --- | --- | --- |
| [2,2] | 15.2s | 74.1% |
| [3,3] | 6.8s | 85.3% |
| [4,4] | 3.9s | 88.9% |
| [5,5] | 2.6s | 95.6% |
| CNN | 20.3s | 95.1% |

**Table 3.** Optimization results of different quantity of features

| Number of features | Running speed | Average accuracy |
| --- | --- | --- |
| 0 | 0.03s | 80.5% |
| 1 | 1.1s | 82.5% |
| 2 | 1.3s | 86.9% |
| 3 | 2.6s | 95.6% |
| 4 | 3.4s | 89.5% |

**Figure 9.**
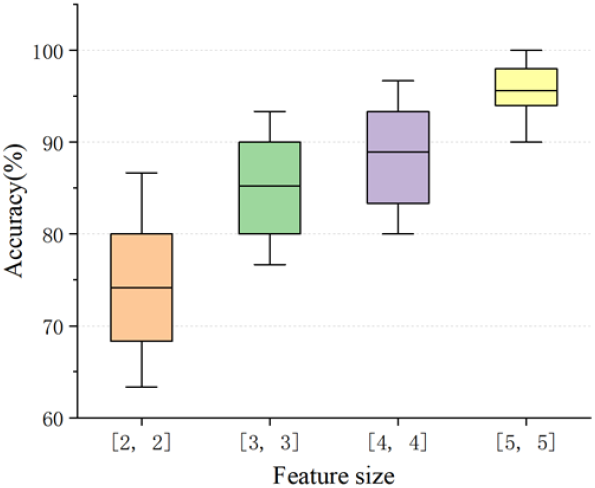
Precision of different window dimension.

**Figure 10.**
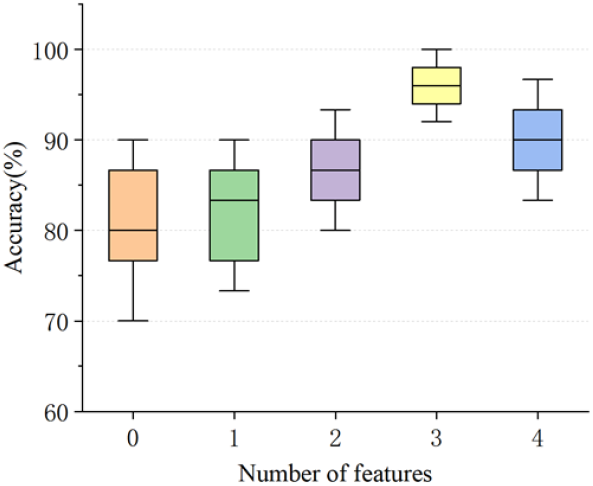
Model precision of different quantity of features.

**Figure 11.**
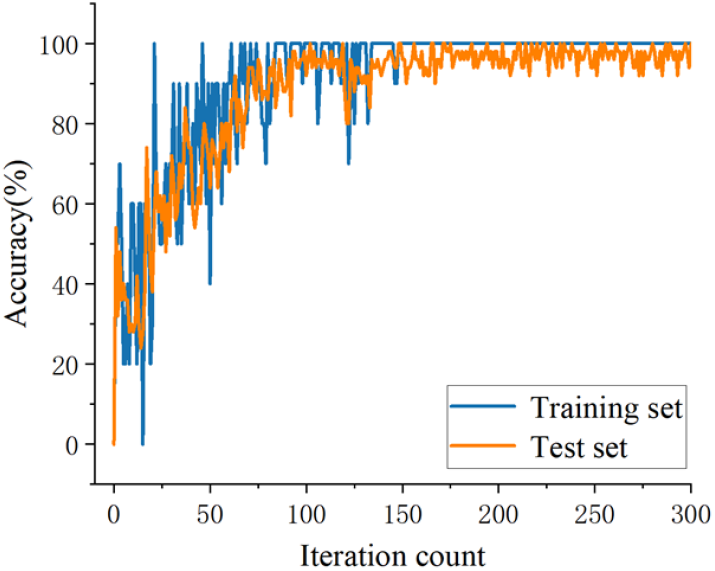
Precision of small window [5,5].

According to Table 2, compared with the classification results of CNN model, PAM-CNN model ran faster, and was more precise when the small window size was [5,5]. According to Table 3, when the small window size was [5,5], the model running speed decreased and its accuracy showed a non-linear trend as the number of features increased. Moreover, when the number of fatigue features was 3, PAM-CNN could approach a significant precision 95.6%.

## 4. Conclusions

1. Oriented to the fatigue problem of special operators, research on the mechanism of muscle fatigue was carried out, a five-element model based on hidden Markov chain was built, feature samples for fatigue recognition were established by feature extraction using time-overlapping sliding window and cmor wavelet on the sEMG data of the operators, and a Fatigue recognition model based on PAM-CNN was proposed.
2. For container crane operators, a case study of the proposed model in this research was introduced, and the cross-entropy-adam algorithm was used to optimize the number of features and the size of small window. The results showed that when the number of features was 3, and the size of small window was [5,5], the average accuracy could reach 95.6%. Compared with the classification results of CNN algorithm, the PAM-CNN model ran faster and had a higher average accuracy.
3. The next step would consider the change in hip muscle pressure during the operator’s work process, furthermore introduce EEG and ECG signals, in order to carry out multimodal information fusion, and establish big data samples to improve robustness of fatigue recognition.

## Data Availability Statement

Some or all data, models, or code that support the findings of this study are available from the corresponding author upon reasonable request.

## Acknowledgement

This paper is sponsored by the National Natural Science Foundation of China (No. 51675345); Science and Technology Commission of Shanghai Municipality Capacity Building Plan for Some Regional Universities and Colleges (20090503000); Biomedical science and technology support project of Science and Technology Commission of Shanghai Municipality (20S31905500); Science and technology talent development fund of SIT(ZQ2020-23); Public welfare projects of the State Administration for Market Regulation (2020MK030, 2024MK030).

## Ethics Statement

This study did not involve ethical problem, and all experiments provided written informed consent.

## Author Contributions Statement

Xu Bin: data organization, software, investigation, writing, review, and editing.

Lin Minghui: data organization, software.

Yue Zhongzhen:writing, and editing. Ji Shoucheng: review.

Ouyang Weiping: writing check. Li Changyu: data process.

## Conflict of interest statement

The authors declare that they have no conflict of interest.

## Notes

### Competing Interest Statement

The authors have declared that no competing interests exist.

